# The Quest to Identify USP8 Inhibitors for Parkinson’s Disease, a PAINful Experience

**DOI:** 10.1101/2023.09.05.556294

**Authors:** Stuart Lang, Fiona Bellany, De Lin, Denise S Barrett, Kieran R. Cartmill, Daniel A. Fletcher, Catrina Kerr, Andrew Plater, Barbara Forte, Beatriz Baragaña, Parul Dixit, Mairi M. Littleson, Mary C. Wheldon, David W. Gray, Fraser Cunningham

## Abstract

Pan Assay INterference compoundS (PAINS) are known to be a source of false positives in High Throughput Screening (HTS) campaigns. This has become a major problem in medicinal chemistry, often resulting in undesirable project outcomes and increased overall cost. Our recent campaign to identify inhibitors of USP8 that could be used in the treatment of Parkinson’s disease identified several PAINS that worked *via* a variety of mechanisms. Herein, we discuss the process developed to identify not only the PAINS but also confirming the interference mechanism causing their activity. We found in this project that our USP8 assay was susceptible to multiple modes of interference, making it difficult to identify genuine hit molecules.

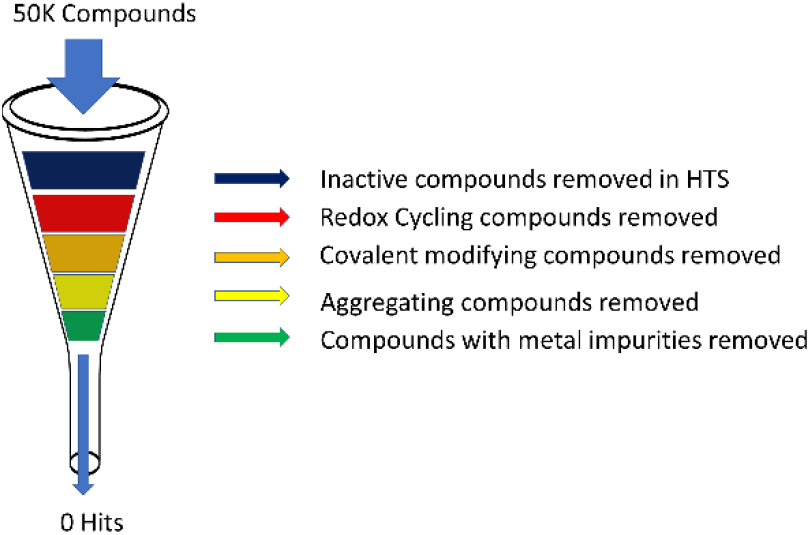

## Introduction

The accumulation of misfolded and often ubiquitinated α-synuclein in neuronal inclusions known as Lewy Bodies is a key pathological marker of Parkinson’s disease. In 2016 Alexopoulou *et al*.,^1^ demonstrated that Lewy Bodies contain predominantly K63-linked ubiquitin chains and that the quantity of these conjugates was inversely corelated with the localisation of Ubiquitin-specific protease 8 (USP8), a deubiquitinating enzyme (DUB) from the Ubiquitin-specific protease family. They proposed that up regulation of USP8 could lead to a reduction in clearance of α-synuclein resulting in its accumulation and the formation of neurotoxic Lewy bodies.

In addition to this, USP8 has been shown to play a role in numerous other biological processes with its effects been seen in areas such as cancer,^2^ inflammation^3^ and male fertility.^4^ To this end it was reasoned that the discovery of novel molecules that can modulate the function of USP8 could have benefits that reach beyond those of Parkinson’s disease.

DUBs fall into two main classes, cysteine proteases and metalloproteases, with the USP family being the largest of the cysteine proteases.^5^ Due to the crucial role that the catalytic cysteine plays in the function of these types of enzymes, we were acutely aware of the problems that assay interference could play in the apparent activity of any inhibitors identified.^6^ The term Pan Assay Interference compoundS (PAINS) was introduced in 2010 by Baell *et al*.^*7*^ to describe molecules that display activity against protein targets that cannot be attributed to a mechanism or interaction that is unique to the protein of interest.^8^ Well documented PAINS mechanisms include aggregation,^9^ covalent modification,^10^ redox cycling^11, 12^ and sample impurities^13-15^. In addition to this the general promiscuity of a compound, resulting in a high Inhibitory Frequency Index (IFI),^16^ is a sign of assay interference even if the specific method of interference has not been identified through triaging and counters-screening assays.

Herein, we describe the application of a biochemical screening platform that was used for the screening of our diverse in-house compound screening collections to identify inhibitors of USP8. Whilst this screen produced potential hit molecules, each compound failed to reconfirm in an orthogonal technique, binding to USP8 by either SPR or NMR, and this led us to further investigate the potential mode of inhibition of these compounds by a panel of different assays to determine if the hits were genuine. This assessment was not limited to their measured potency, but also to understanding if this potency could be attributed to a desired specific interaction with USP8 or if it was because of a non-specific interference mechanism. Surprisingly the compounds disclosed in this paper, that have shown activity against USP8, have all been shown to be active through non-specific mechanisms and their potency cannot be attributed to genuine activity.

## Results

### Screening Platform

The biochemical screening platform used for our diversity screen was based upon RapidFire mass spectrometry (RapidFire-MS) technology, which conducts solid phase extraction and allows for an accurate quantitation of enzyme reaction product. This screening platform has been successfully used by us and others for library screening in many targets including SARS-CoV-2 guanine-N7-methyltransferase,^17^ leucyl aminopeptidase,^18^ acetyl-coenzyme A synthetase^19^ and S-adenosylmethionine decarboxylase.^20^

For our diversity screen, we developed a RapidFire-MS based assay to assess the activity of USP8 through the detection of Ac-KKTIPNDSRE, the de-ubiquitinated product of our model substrate (Ac-KK(Ub)TIPNDSRE). This enabled us to screen 50k compounds from our in-house libraries to identify potential inhibitors of USP8.

### Literature DUBs Inhibitors

In addition to identifying potential starting-points for our drug discovery program through screening, we also searched the literature for inhibitors of the different members of the USP family (Figure 1). ^5, 21, 22^ The compounds chosen were reported to have activity against various USPs, with a bias toward compounds that had shown activity against DUBs that were closer to USP8 in the phylogenetic tree.^5^ Unfortunately this exercise did not identify any molecules with measurable activity against USP8. While this was disappointing, it gave us confidence that selective inhibitors could be developed.

**Figure 1.**
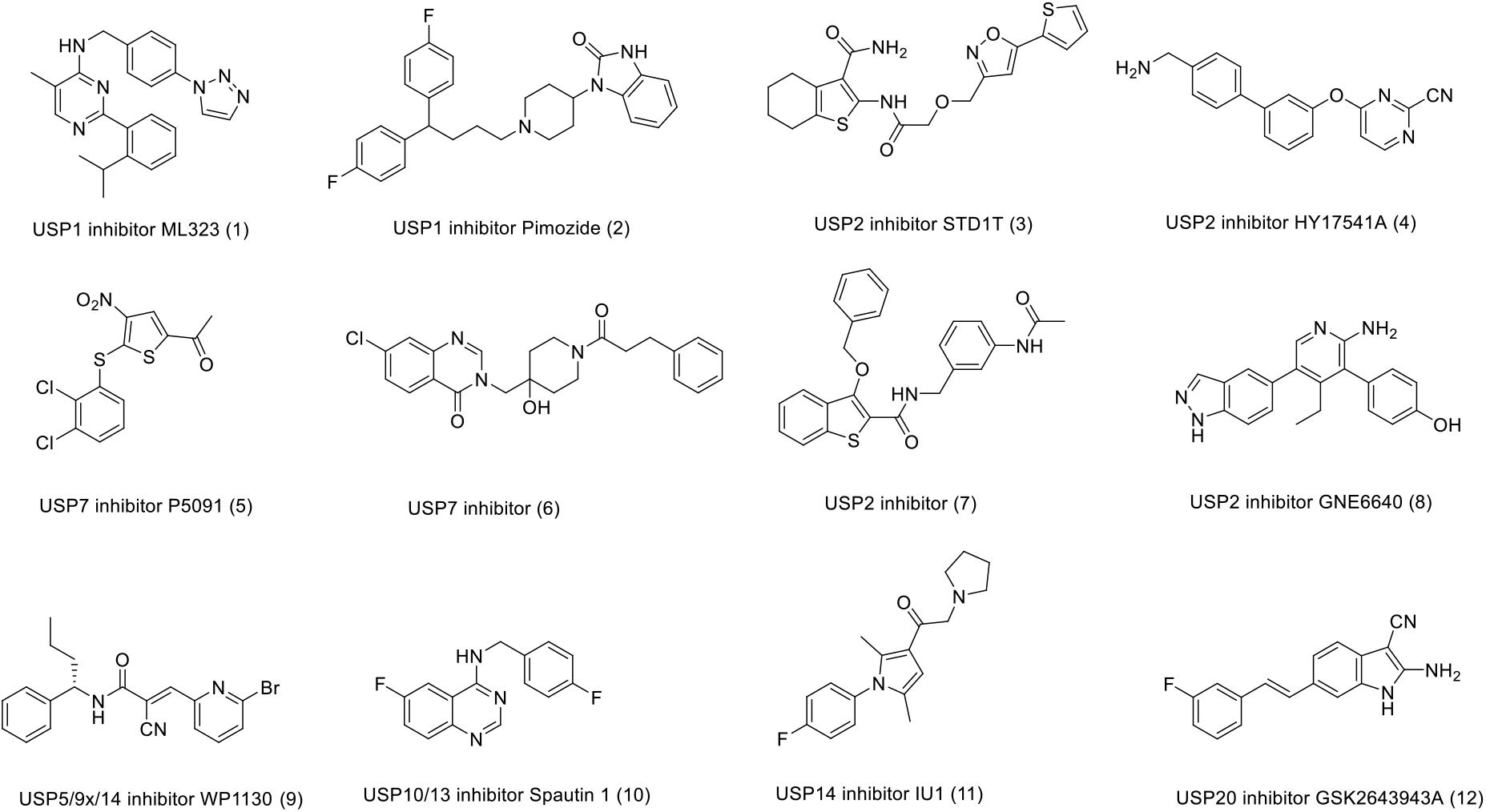
Published USP inhibitors that were inactive when tested against USP8^5, 21^

**Scheme S1.**
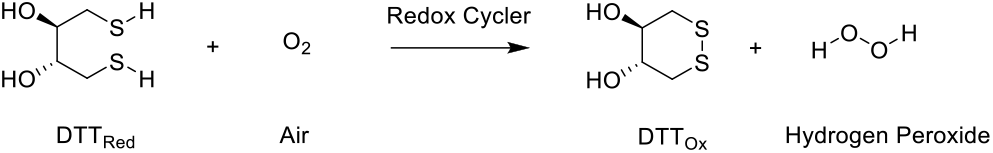
DTT mediated production of hydrogen peroxide from air.

### Redox Cycling

There are several examples of molecules that have been reported to be pan DUB inhibitors, many of these compounds are known to be redox cycling compounds (RCC).^23^ A common trait of enzyme targets, such as DUBs, affected by redox is the catalytic cysteine residue in the active site. The oxidation of the thiol residue prevents further catalytic turnover of the enzyme, rendering it inactive.

DTT is known to facilitate the production of hydrogen peroxide from oxygen in the atmosphere.^12^ While real, this process is very slow and without significant catalysis does not interfere with assays. The addition of a RCC can catalyse this process meaning that it only takes minutes to generate enough hydrogen peroxide, a strong oxidising agent, that can result in assay interference (Scheme 1). The sensitivity of USP8 to inhibition by hydrogen peroxide was measured against USP8 and found to have a pIC_50_ of 5.7 in our assay.

By adapting the horseradish peroxidase (HRP) assay that oxidises phenol red in the presence of H_2_O_2_ developed by Johnson *et al*.^*24*^ we were able to identify compounds that had the ability to redox cycle. An important note regarding this assay; we found its sensitivity was only suitable for identifying compounds producing greater than 30 µM of H_2_O_2_. Lower than this and the signal found on our plate reader was too close to background to confidently identify a positive RCC. Importantly the lower limit of quantification of the HRP assay was 15 times less than the IC_50_ of peroxide in our assay. Interestingly we also observed these high levels of sensitivity to oxidation by hydrogen peroxide against other proteins we’ve recently worked on at the Drug Discovery Unit, such as the related DUB USP15 and the unrelated *Plasmodium falciparum* Serine tRNA synthetase (*Pf*SerRS). To understand if the HRP redox assay is suitable to identify RCCs that will interfere with an assay we recommend including a titration of hydrogen peroxide against any potential protein targets.

When working with redox sensitive proteins one key factor is the choice of reducing agent. As described above the reducing agent used in an assay such as DTT can catalyse the production of hydrogen peroxide. Replacement of these catalytic reducing agents with a weaker (non-catalysing) reducing agent such as glutathione (GSH) significantly reduces the rate that hydrogen peroxide is produced.^12^ This means that the use of GSH is less likely to make compounds appear active because of redox cycling. Under these ‘non-redox cycling conditions’, most compounds which were identified as RCCs in the HRP assay had significantly lower pIC_50_s against USP8.

Whilst conducting our search for literature DUBs inhibitors, many compounds identified contained potential redox motifs (Table 1), so we sought to validate their activity using the various screening conditions we had developed: our USP8 assay using either DTT or GSH as the reducing agent and our HRP redox assay. These compounds all displayed a pIC_50_ between 4.9 and 6.6 against USP8 using DTT as the reducing agent. Whilst most of these compounds displayed significant activity in our HRP assay, above a pIC_50_ of 5.0, there were some examples that were not obvious. Entry 3 displayed only moderate activity and entry 4 showed no activity at all, despite both containing an embedded 1,4 quinone scaffold within the core of the molecule. In addition to these two molecules entry 6 also displayed no redox activity in our HRP redox assay. All molecules however have significantly lower activity when GSH is used as the reducing agent in place of DTT. This highlights the increased sensitivity in detecting redox cycling when comparing the difference in activity when moving from using DTT to GSH as a reducing agent as opposed to purely measuring redox activity in a HRP assay. This lower activity when GSH is used suggests that all or some of the effects of these compounds can be attributed to their ability to generated H_2_O_2_ under the assay conditions.

**Table 1.**
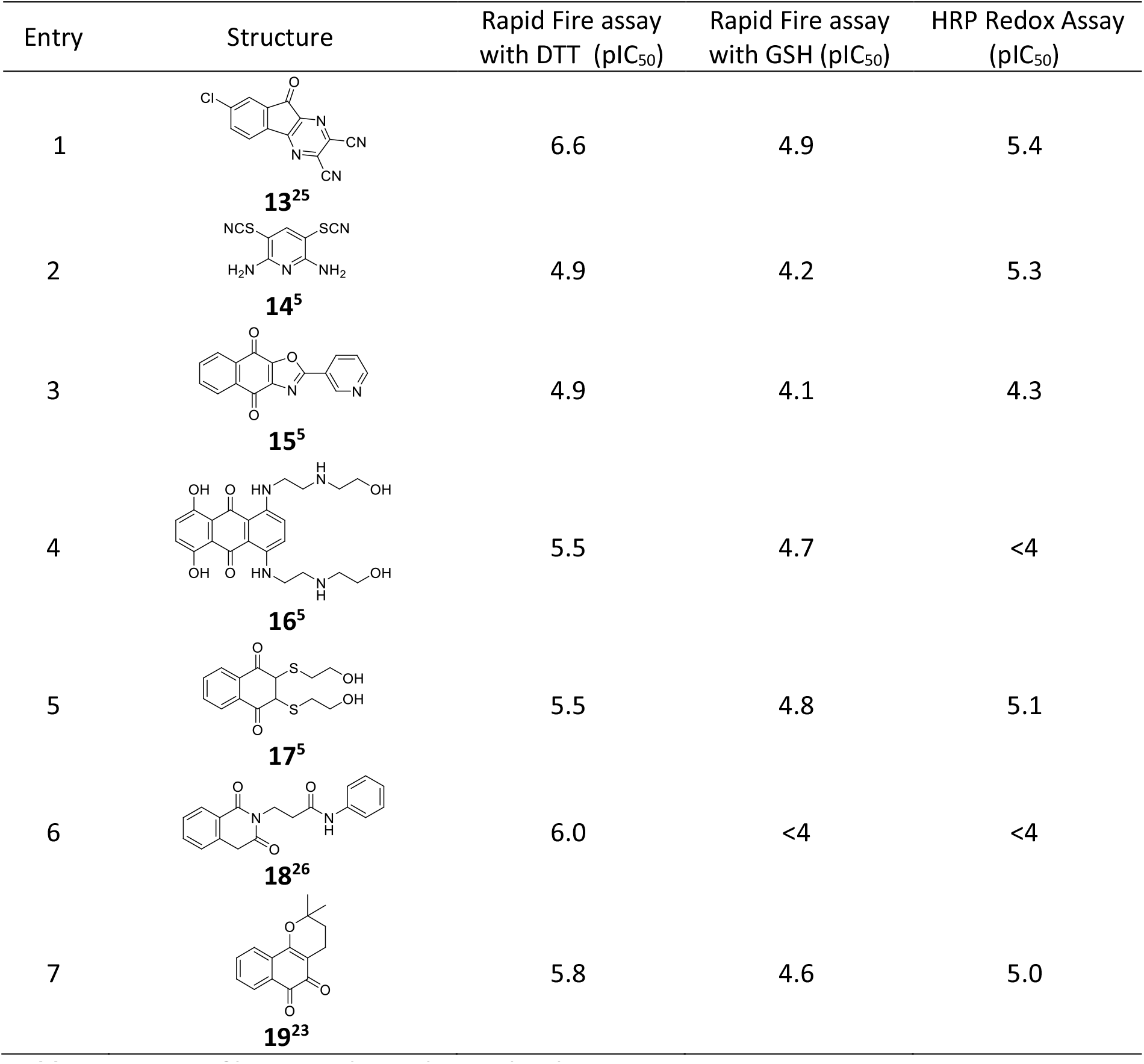
Potency of known redox cycling molecules against USP8.

| Entry | Structure | Rapid Fire assay<br>with DTT (pIC<sub>50</sub>) | Rapid Fire assay<br>with GSH (pIC<sub>50</sub>) | HRP Redox Assay<br>(pIC<sub>50</sub>) |
| --- | --- | --- | --- | --- |
| 1 | **13<sup>25</sup>** | 6.6 | 4.9 | 5.4 |
| 2 | **14<sup>5</sup>** | 4.9 | 4.2 | 5.3 |
| 3 | **15<sup>5</sup>** | 4.9 | 4.1 | 4.3 |
| 4 | **16<sup>5</sup>** | 5.5 | 4.7 | <4 |
| 5 | **17<sup>5</sup>** | 5.5 | 4.8 | 5.1 |
| 6 | **18<sup>26</sup>** | 6.0 | <4 | <4 |
| 7 | **19<sup>23</sup>** | 5.8 | 4.6 | 5.0 |

Many screening campaigns carried out within the DDU have shown the compounds in Table 2 to be active, as illustrated by their high IFI^16^ scores, with the compounds being active in 17-21% of screens. This promiscuity has caused compounds of this type to be treated with caution when appearing active in an in-house diversity screen. Our USP8 screen identified these compounds as hits when screened using DTT as the additive. When the DTT is replaced with GSH the activity drops, which correlated with their activity in our HRP redox assay and confirms that these compounds do indeed redox cycle. Furthermore, these compounds are not stable in solution and oxidise to the give a fully aromatic central pyridone ring. These known assay interference mechanisms provided some rationale as to why the molecules appear active in so many screens and that their activity is not suitable for developing into drugs.

**Table 2.**
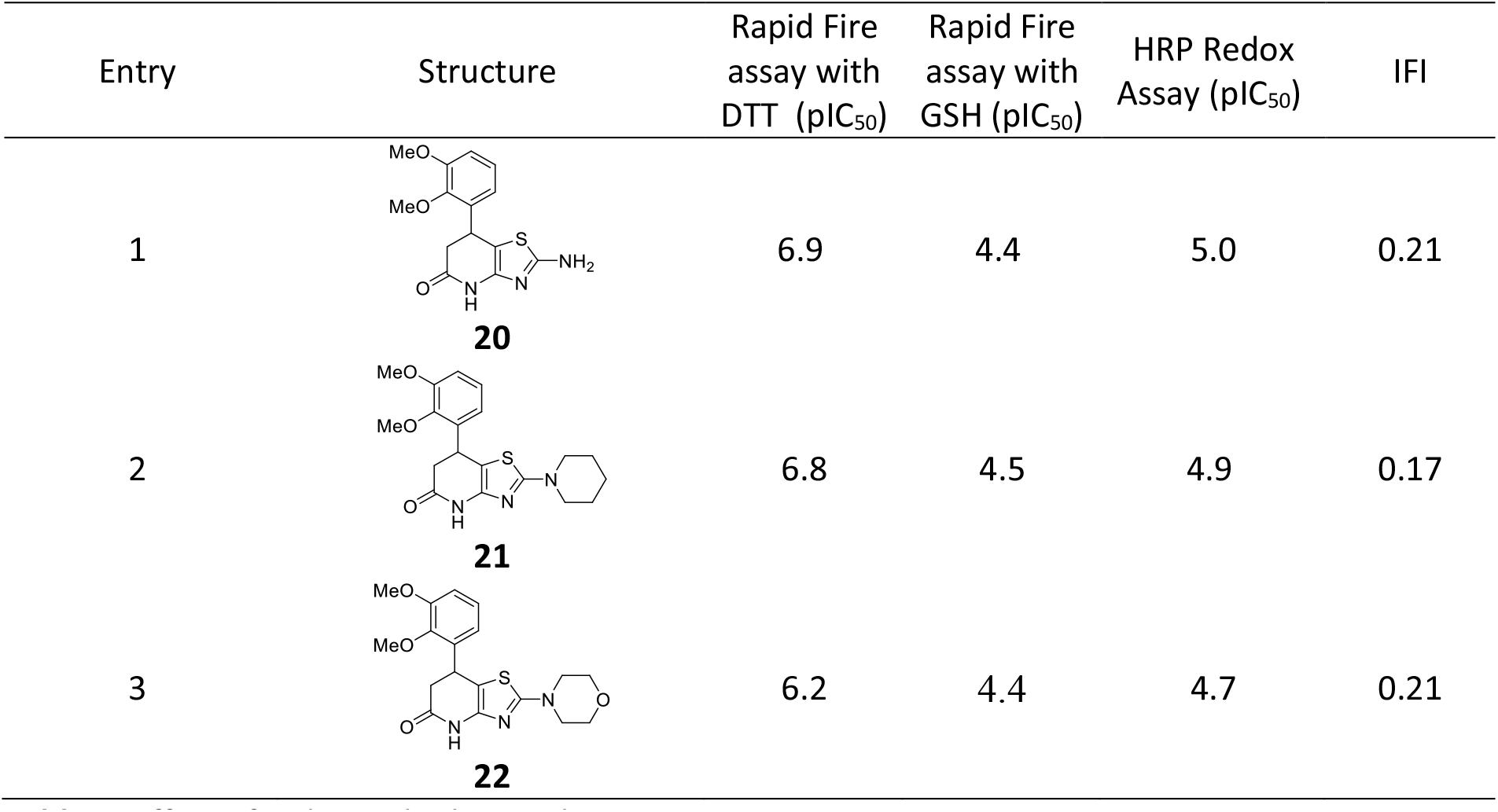
Effect of redox with observed potency.

| Entry | Structure | Rapid Fire assay with DTT (pIC<sub>50</sub>) | Rapid Fire assay with GSH (pIC<sub>50</sub>) | HRP Redox Assay (pIC<sub>50</sub>) | IFI |
| --- | --- | --- | --- | --- | --- |
| 1 | **20** | 6.9 | 4.4 | 5.0 | 0.21 |
| 2 | **21** | 6.8 | 4.5 | 4.9 | 0.17 |
| 3 | **22** | 6.2 | 4.4 | 4.7 | 0.21 |

The effect of redox potential on potency was observed with a second series of molecules identified from our diversity screen against USP8 (Table 3). Initial tests of the hit chemical matter in the HRP redox assay during hit validation indicated that these molecules were not RCCs and initially these molecules appeared to have good potency along with high ligand efficiency. However, after a short round of optimisation of this series and, with more potent analogues prepared, their activity in the redox assay was checked again and unfortunately it showed that as the potency in the DTT USP8 assay was increased so did the activity in the redox assay. Effectively this showed that the SAR of this series was being purely driven by redox activity, which initially was not strong enough to appear active in the redox assay but was strong enough to inhibit USP8, highlighting the exquisite sensitivity of this enzyme to RCCs.

**Table 3.**
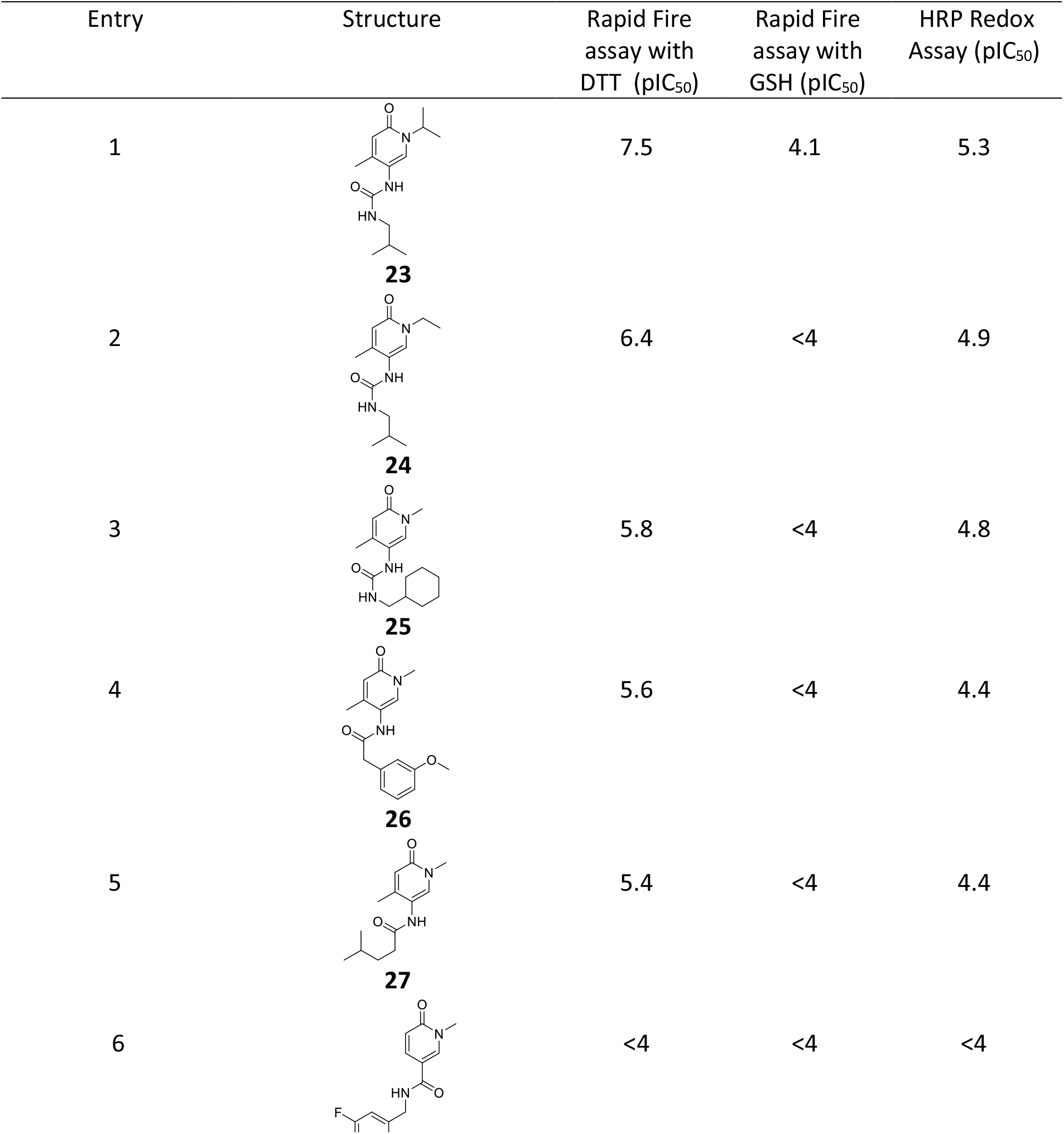
Effect of redox with observed potency.

| Entry | Structure | Rapid Fire assay with DTT (pIC<sub>50</sub>) | Rapid Fire assay with GSH (pIC<sub>50</sub>) | HRP Redox Assay (pIC<sub>50</sub>) |
| --- | --- | --- | --- | --- |
| 1 | **23** | 7.5 | 4.1 | 5.3 |
| 2 | **24** | 6.4 | <4 | 4.9 |
| 3 | **25** | 5.8 | <4 | 4.8 |
| 4 | **26** | 5.6 | <4 | 4.4 |
| 5 | **27** | 5.4 | <4 | 4.4 |
| 6 | **28** | <4 | <4 | <4 |

In a similar manner to the known literature examples, replacing DTT with GSH in the assay conditions resulted in a reduced activity with pIC_50_ values below or equal to 4.1 in all cases with most being below the detection limit of the assay. These compounds also displayed measurable levels of activity in our HRP redox assay which further confirmed that redox, driven by the embedded azo-quinone core, was the mechanism that allowed these compounds to inhibit USP8. It is noteworthy that reversing the amide substituent (Table 3, Compound **28**), making the compound unable to undergo redox cycling, rendered this molecule inactive in all our assays.

The observation of the link between redox and USP8 activity, which appears to be purely driven by the production of hydrogen peroxide made it clear that it would not be possible to optimise any molecule that displayed redox activity in our assay with DTT as the reducing agent. In all cases replacement of DTT with GSH resulted in a loss of activity so it is unlikely that the activity of any of these molecules was due to a specific interaction of the molecule with USP8.

This series was also identified in an unrelated target to USP8 and was identified in a project looking for inhibitors of Serine tRNA synthetase (*Pf*SerRS). *Pf*SerRS belongs to the tRNA synthetase family. This family of enzymes catalyses the attachment of amino acids to their cognate tRNAs to produce the aminoacyl tRNAs that are the substrates for translation. This family are generally druggable targets. Compounds that inhibit aaRSs have been successfully exploited, with at least one antibacterial drug, mupirocin, currently in clinical use for the topical treatment of Staphylococcus aureus, that acts through the inhibition of the isoleucyl-tRNA synthetase (IleRS) of gram-positive bacteria. ^27^ A series of other specific aaRS inhibitors have been also developed,^28^ some of which are currently in clinical trials as antimicrobials. ^29^

An initial diversity screening campaign to identify inhibitors of *Pf*SerRS was conducted with assay conditions using DTT as reducing agent and led to the identification of several false positive hits which were operating *via* a redox-cyclising interference mechanism, including those identified in Table 3 (see supporting information). Compounds were assessed in NMR binding experiments and proved not to bind to PfSerRS. A cysteine is present in proximity of PfSerRS ATP binding site (Cys 345) and modelling studies suggested that oxidation of this Cys residue by the H_2_O_2_ generated in the assay conditions could prevent ATP from binding to the active site. To remove the risk of identifying further redox cycling compounds, the assay conditions were adapted to use 2-mercaptoethanol (a weaker reducing agent) and this resulting in a loss of activity for the redox cycling compounds.

### Covalent modifier

Compound **29** is a published PAN DUB inhibitor (Figure 2).^30^ Using our GSH assay conditions, that reduce the effect of redox cycling, it was found to be active with a pIC_50_ of 5.1 against USP8. While this molecule does not appear to be a redox active molecule, the α-chloro ketone moiety is known to covalently bind to amino-acids such as serine and cysteine. While selective covalent interaction with the catalytic cysteine could be desirable, what was less desirable is a molecule where the covalent reactivity is un-selective. As part of our efforts to understand the importance of the chlorine atom that gives rise to the reactivity, the non-covalent analogue **30** was also assessed for comparison.

**Figure 2.**
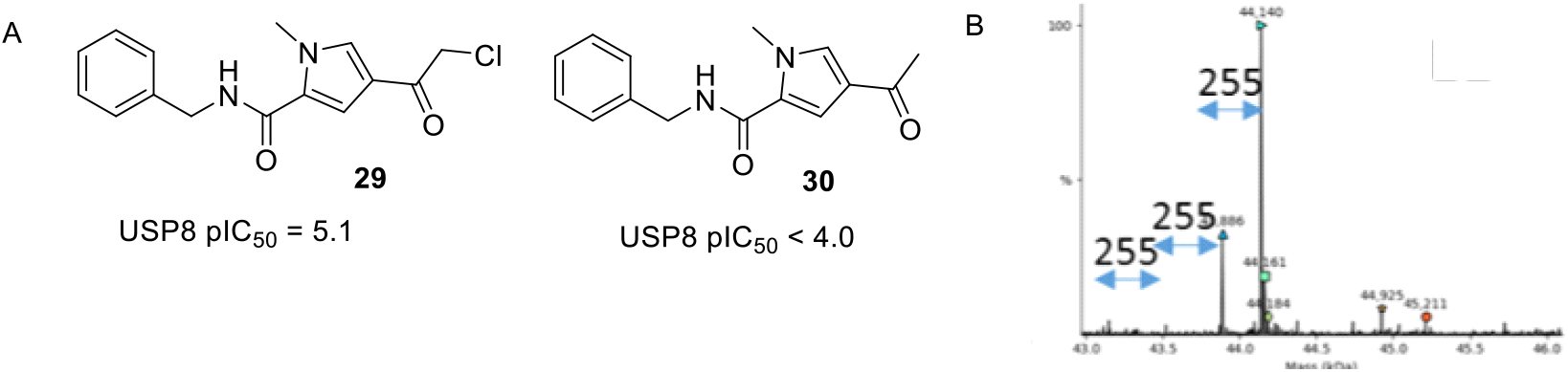
Covalent Pan DUB inhibitor along with its non-covalent analogue.

**Figure 3.**
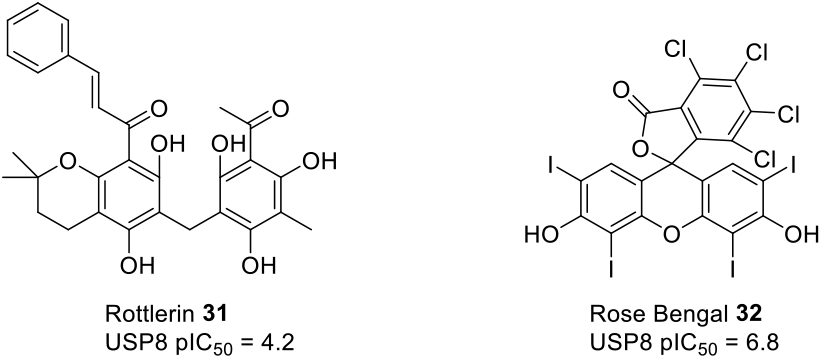
Compounds reported in literature to be aggregators that show activity in USP8 assay.

The lack of activity of the **30** highlights the importance of covalent reactivity in the observed potency in **29**. To better understand the specificity of this compound we used intact mass spec analysis to identify the stoichiometry of the covalently modified protein (Figure 2B). The most common adduct contained 3 equivalents of ligand compound with very few products with only 1 addition observed. This shows that this molecule not only didn’t have selectivity toward a specific DUB but that it has no selectivity towards a specific residue within USP8. This indicates that the molecule is a promiscuous binder with little or no specific recognition through non-covalent interaction with USP8. This means that the covalent warhead in this molecule is acting as the main source of activity rather than providing an irreversible binding interaction that is primarily driven by a genuine non-covalent interaction with the target.

### Aggregation

It is known molecules that aggregate can form large polymolecular assemblies that could sequester and denature proteins therefore giving a similar effect to that of inhibition. These aggregates can contain as many as 10^4^ molecules as opposed to the single molecule that would be required for a genuine inhibitor.^31^ As this effect is not due to a desired interaction of the molecule with the protein, molecules that form aggregates often display promiscuous activity against several unrelated protein classes, with analogues within a series having little or no discernible SAR. This in turn leads to an optimisation cul-de-sac with little improvement seen after multiple design iterations.

As part of our exercise to profile known USP inhibitors against USP8 we found that Rottlerin, a published USP1 inhibitor displayed a weak pIC_50_ of 4.2 in our GSH-USP8 assay. Rottlerin is also known to be an aggregator,^32^ this led us to suspect that our USP8 assay may be susceptible to identifying molecules that can form aggregates and displaying them as false positives. This was supported further when Rose Bengal, another known aggregator^9, 32^ with no prior evidence of USP8 activity, appeared to have a pIC_50_ of 6.8.

It is common practice to determine if compounds are aggregating under assay conditions by running the assay in the presence and absence of a non-ionic detergent, such as Triton X-100 or Tween-80. The addition of these detergents serves to disrupt aggregates and therefore the appearance of potency in their absence. Potency being lost when detergent is added is a sign that the compounds are aggregating under the assay conditions. This method of identifying aggregation was not possible in our USP8 assay because high concentrations of detergent were incompatible with the MS platform that is used.

To determine whether a compound displays aggregating behaviour, we used the proton NMR methodology developed by LaPlante *et al*.,^33^ where compounds are screened at varying concentration in buffer (pH8 20mM Tris 150mM NaCl 1mM TCEP 10% D2O) and the chemical shift at these concentrations compared. Compounds that display aggregation would result in a drift in the chemical shift, whereas compounds that do not aggregate would show no change in chemical shift across all concentrations.

Through our diversity screen we identified a quinoxalinone series (Figure 4) as having potential USP8 activity. To assess this series, we looked at assessing compounds with small point changes that should lead to improved solubility and hopefully reduced aggregation. It was hoped that substitution of the cyclic NH (compound **33**) would result in reduction of the intermolecular hydrogen bonding within the molecule. The addition of methyl groups to the aromatic ring either with two Me groups added (compounds **33** and **34**) or a single Me group added (compound **36**) was expected to break any planarity that exists within the molecule, by increasing the overall fraction of sp^3^ character of the molecule. Additional known solubilising groups, such as morpholine (compounds **34**) and methyl ether (compound **35**) were also tested to improve the solubility of these molecules. However, all these modifications gave negligible changes in potency and all these molecules were subsequently shown to be aggregators in the previously described NMR aggregation assay, despite also displaying good solubility (>150µM).

**Figure 4.**
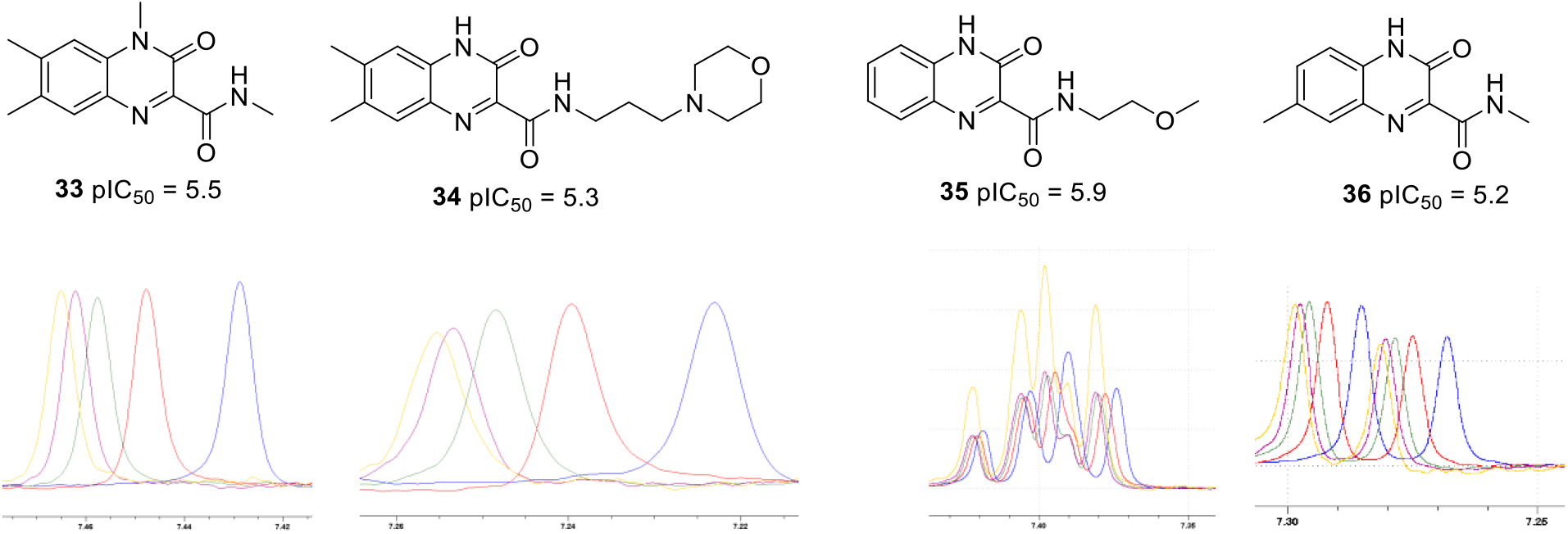
NMRs carried out at various concentrations with the following colour coding: 200µM (Blue), 100µM (Red), 50µM (green), 25µM (purple), 12.5 µM (yellow).

**Figure 5.**
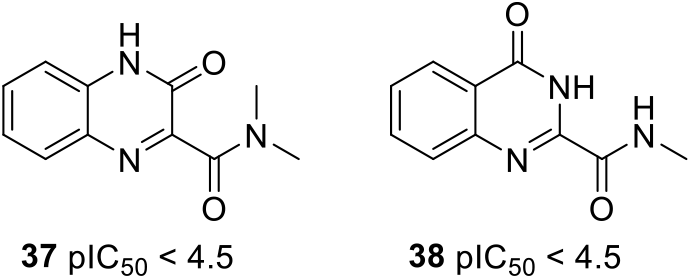
Inactive compounds.

This aggregation could be due to hydrogen bond that exists between the exocyclic amide N-H and the carbonyl within the ring. Substitution of the amide N-H, meaning that intramolecular hydrogen bonding is no longer possible, resulted in a complete loss in activity (compound **37**) as does reversing the amide substituents in the ring (compound **38**). This highlights the importance of testing for aggregation when carrying out the HTS process. Our inability to carry out routine experiments using detergents meant that we required to use this alternative NMR technique to confirm that these hits were not genuine inhibitors, but instead were displaying the appearance of inhibition purely because of compound aggregation.

### Metal Impurities

Assay interference is also known to result from metal impurities contained within in the sample.^13^ These impurities are often an artefact of the synthetic route used to prepare the compound, which may have used transition metal catalyst, with the subsequent purification process not adequately removing these trace metals. As compounds are routinely analysed using methods such as NMR and LCMS, which only detect organic compounds, impurities of this type can enter compound screening collections without being detected. This problem can be particularly prevalent when the molecule being prepared contains multiple heteroatoms, such as nitrogen atoms. This allows the molecule to chelate the metal ion meaning removal can be more problematic.

Even when present in trace quantities, these metal impurities are known to make compounds appear active in assays. Even although the testing the molecule in the absence of the metal, for example after the sample has been treated with a metal scavenger resin, no activity is observed. This activity can also be due to previously discussed redox methods. Only this time it is the metal or the metal-compound complex that causes the redox activity rather than the molecule alone. It can also be due to the protein being particularly sensitive to metal ions, which can lead to denaturing on the protein leading to a perceived reduction in activity of the protein.

Using our previously described assay condition containing DTT as an additive, which is known to produce hydrogen peroxide from atmospheric oxygen, compounds **39** and **40** (Table 4) appeared to be active with a pIC_50_ of 5.2 and 5.6 respectively. When these compounds were subsequently tested using the GSH, as a weaker reducing agent, both compounds were inactive in these modified assay conditions. This result is indicative of the activity of these samples being due to hydrogen peroxide formation coming from the redox activity of the compound. Unlike the previous compounds (Tables 1-3) that were shown to display redox activity even after scavenging of the samples with a MP-TMT resin^34^, which is designed to remove metals such as copper, iron, nickel, palladium, platinum, silver and zinc. The activity of compounds **39** and **40** was lost after scavenging of the samples.

**Table 4.**
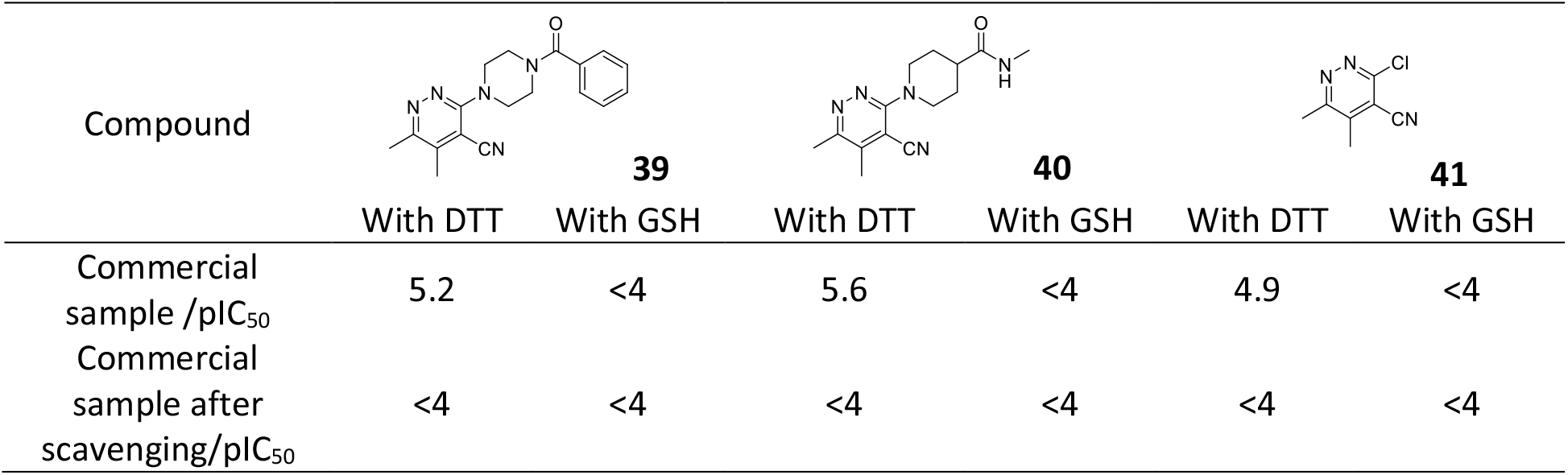
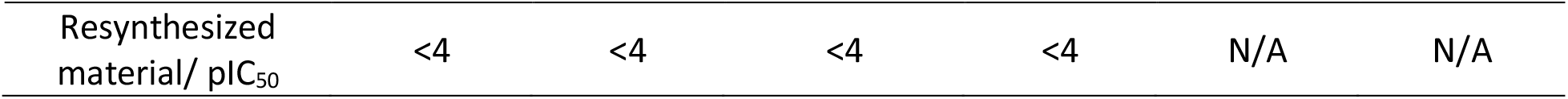
Effect of Metal Impurities on Redox Cycling.

| Compound | **39** |  | **40** |  | **41** |  |
| --- | --- | --- | --- | --- | --- | --- |
|  | With DTT | With GSH | With DTT | With GSH | With DTT | With GSH |
| Commercial sample / $pIC_{50}$ | 5.2 | <4 | 5.6 | <4 | 4.9 | <4 |
| Commercial sample after scavenging/ $pIC_{50}$ | <4 | <4 | <4 | <4 | <4 | <4 |
| Resynthesized material/ pIC<sub>50</sub> | <4 | <4 | <4 | <4 | N/A | N/A |

Furthermore, in-house resynthesis of these compounds resulted in samples that were inactive in our USP8 assay. This further suggested that all activity displayed with these compounds was due to redox activity brought about, not by the compound alone but, a metal impurity or the complex of the metal and the compound. Testing of the starting material **41** used to prepare compounds **39** and **40** also displayed activity in the assay conditions where DTT is added as an additive but not the conditions where GSH is added as an additive. This is further evidence that the metal impurity was present in the common starting material used to make compounds **39** and **40**.

Compound **42** and **43** appeared active against USP8 when GSH was used as an additive (Figure 6). This indicates that the activity of these samples was not due to a redox mechanism. However, scavenging of these compounds with the MP-TMT resin resulted in a loss of activity meaning that the molecules themselves were not active and that the activity was due either the metal contaminant or a complex with the molecule and the metal contaminant. Elemental analysis of this sample showed that the original sample of **42** contained 5.5% of copper, which was successfully removed by the scavenger resin indicating that our USP8 assay is sensitive to copper impurities within the samples.

**Figure 6.**
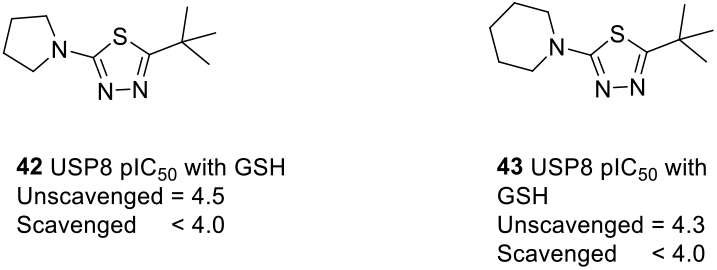
Contaminated compounds that do not show redox activity.

These results highlight that while metal impurities have caused the interference in both series this interference has shown itself in different ways. In the example of **39** and **40** the metal that was present in the starting material caused the formation of hydrogen peroxide *via* a redox reaction. This effect of this interference could be eliminated by using GSH as an additive instead of DTT, however DTT could still be used if the metal was removed. However, in the example of **42** and **43** the effect of the metal impurity, later identified as copper, did not cause the formation of hydrogen peroxide *via* a redox reaction. This impurity was causing interference *via* an alternative mechanism that did not take place when the copper impurity is absent from the sample.

As a result of these finding we recommend that all hits are synthesised and purified to confirm activity. If this is not possible due to resource or time constraints, then we recommend that compounds sourced from commercial supplies are purified using metal scavenging technology prior to confirming activity.

## Conclusion

While we were unsuccessful in identifying a genuine inhibitor for USP8, this doesn’t mean that identifying such compounds is not possible. By screening alternative compound collections, the identification of active compounds whose activity is not due to interference may still be possible. We have shown that USP8 is a particularly sensitive protein to numerous forms of assay interference. Therefore, we recommend that any exercise aimed at identifying a novel USP8 inhibitor requires a robust suite of assays and counter screens to rule out the numerous modes of interference as the source of apparent potency.

We have demonstrated that understanding how sensitive a protein is to hydrogen peroxide is helpful when initiating screening as the sensitivity of the industry standard HRP assay can be lower than that of target proteins. We therefore recommend that if there are concerns that a target protein is sensitive to RCCs then a full titration with hydrogen peroxide should be carried out to determine the level of sensitivity to this source of interference. Ideally screening should be carried out with weaker/ non-catalysing reducing agents such as GSH but if this is not possible the HRP redox assay should be used to counter screen hits.

Prior to beginning this campaign, we were aware of the issues that could be experienced because of the catalytic cysteine present in proteases such as USP8. Not only could this moiety be oxidised because of redox, rendering the enzyme inactive, but it could also irreversibly bind to reactive covalent warheads. Using intact mass spec, we were able to show that molecules can covalently bind multiple times to USP8. This indicates that the catalytic cysteine in the active site is not the only reactive cysteine present in USP8. This means that the development of inhibitors with covalent warheads for targeting USP8 should be treated with extreme caution as the protein-inhibitor stoichiometry could be difficult to control.

In addition to these predicted challenges associated with USP8 we have also shown that this protein is sensitive to factors such as aggregation and metal contamination of samples. Both compounds reported to aggregate in the literature and compounds we identified that were subsequently shown to aggregate appeared active against USP8. Unfortunately, due to the screening technology being used we were unable to modify the detergent concentration to control for aggregators, but an alternative NMR method was able to confirm the aggregation of these compounds.

Metal impurities, which can be present in samples to be tested in biological assays can also lead to USP8 false positives, with these both displaying redox and non-redox modes of activity. We recommend that all hits should be resynthesised and purified to confirm activity but if this is not possible then compounds from commercial sources should be purified with metal scavenging technology before confirming activity.

This paper outlines the processes that can be taken to minimise the impact assay interference can have on a project when working with a sensitive protein, like USP8. While this story has been specific to the challenges faced while working on this protein, we feel that processes and overall strategy should be applicable to all hit identification drug discovery projects.

## Acknowledgement

We gratefully acknowledge funding from MRC Confidence in Concept award (MC_PC_16042) and from Bukwang Pharmaceutical. This work would not have been possible without the input from Darren Edwards and our excellent compound (Alex Cookson, Kirsty Cookson, & Desiree Zeller) and data management teams (James Burkinshaw & Kashish Sharma).

## Experimental Section

### USP8 RapidFire-MRM High-throughput Assay

The enzyme reaction was performed in an assay buffer (50mM HEPES, pH7.5, 1mM GSH or 1mM DTT, and 0.005% Nonidet P40) in a clear F-bottom 384-well polypropylene plate (Greiner, cat. 781101). Briefly, enzyme and substrate Ac-KK(Ub)TIPNDSRE (Cambridge Peptides, Birmingham, UK) were incubated to allow the enzyme reaction to take place. In the assay, 10 nM *Hs*USP8 was incubated with 5 µM Ac-KK(Ub)TIPNDSRE in a final volume of 10 µL. After 60 minutes incubation at room temperature, the reaction was quenched by adding 70 µL 1% formic acid (VWR, Radnor, PA) containing 0.1 µg/mL isotope-labelled peptide Ac-KKTIPNDSR*E (Cambridge Peptides, Birmingham, UK) as internal standard. The sample was directly analysed by RapidFire-MRM.

RapidFire-MRM was performed using a RapidFire 365 system (Agilent, Santa Clara, CA) coupled to a triple quadrupole mass spectrometer 6740 (Agilent). The samples were loaded onto a C18 Type SPE cartridge (Agilent) using deionized water containing 0.1% trifluoracetic [TFA] at flow rate of 1.5 mL/min and eluted to the mass spectrometer using acetonitrile/deionized water (95/5, v/v) containing 0.1% TFA in at a flow rate of 1.25 mL/min. The sipper was washed to minimize carryover with deionized water containing 0.1% TFA followed by acetonitrile/deionized water (95/5, v/v) containing 0.1% TFA. Aspiration time, load/wash time, elution time and re-equilibration time were set to 600, 3000, 5000 and 500ms, respectively, with a cycle time of approximately 10 s. The triple-quadrupole mass spectrometer with electro-spray ion source was operated in positive multiple reaction monitoring (MRM) mode. The detailed setting for the mass spectrometer parameters was as follows: capillary voltage: 3000 V; gas temperature: 350 °C; gas flow: 7 l/min; Nebulizer: 40 psi; sheath gas temperature 300 °C shealth gas flow: 11 l/min and nozzle voltage 1500 V. The MRM transitions (Q1 and Q3) for Ac-KKTIPNDSRE as a reaction product and Ac-KKTIPNDSR*E as an internal standard were set as 614.9/129.1 and 620.4/129.1, respectively. The mass resolution window for both parental and daughter ions was set at as unit (0.7 Da). The Dwell time, fragmentor and collision energy for each transition were 50 ms, 100 v and 8 ev, respectively. Peak areas were integrated, and area ratios of the Ac-KKTIPNDSRE peptide to the internal standard Ac-KKTIPNDSR*E peptide were used for quantitation.

For compound testing, an ECHO 550 acoustic liquid-handling system (LabCyte), was used to dispense testing compound DMSO stock into assay plate, giving a final assay top concentration of 100 µM and 1% DMSO. DMSO was added to both 0% inhibition control wells and 100% inhibition control wells (Non enzyme incubation). Assays were performed by adding 5 µL buffer with *Hs*USP8 protein to all wells except buffer only for 100% inhibition control wells in assay plates before the reaction was initiated with the addition of a 5 µL substrate mix containing substrate Ac-KK(Ub)TIPNDSRE to all wells. The plates were incubated in the plate shaker at 300 rpm at ambient temperature for 60 min, followed by addition of 70 µL of 1% formic acid containing 0.1 µg/mL. The reaction mixture was subjected to RapidFire-MS/MS analysis. The peak area ratio, which is the reaction product (Ac-KK(Ub)TIPNDSRE) divided by its internal standard (Ac-KK(Ub)TIPNDSR*E) was used to calculate the inhibitory activity of the tested compounds. We defined the peak area ratio of the reaction without enzyme as 100% inhibitory activity and that of the complete reaction mixture as 0% inhibitory activity.

ActivityBase XE version 9.2.0.106 from IDBS was used for curve fitting and calculations of IC50 value. All IC50 curves were fit to a four-parameter logistics dose–response regression and IC50 values are presented with 95% confidence intervals with the associated n values.

### USP8 HRP Redox Assay

This assay was adapted from Johnston et al.^23^ The reaction was performed in an assay buffer (40 mM Tris-HCl, pH8.0, 0.2 mg/mL BSA and dH20) in a clear F-bottom 384-well polypropylene plate (Greiner, cat. 781101). For wells containing DTT, a final assay concentration of 0.5 mM of DTT was added to this assay buffer. For wells containing glutathione, a final assay concentration of 2 mM of glutathione was added to the assay buffer. Detection reagent added to all wells was the assay buffer with 800 µg/mL phenol red and 60 µg/mL Horseradish Peroxidase (HRP). Briefly, buffers were made on the day of the assay. Compounds were plated in triplicate. Plate 1 contained just assay buffer, plate 2 contained DTT in the assay buffer, and plate 3 contained glutathione in the assay buffer. 20 µL of buffer with or without reducing agent, depending on plate, was added to all wells, except for 100% effect control wells which contained 200 µM H_2_0_2_. 20 µL of assay buffer containing no DTT was added to the 100% control wells in all plates. The plates were centrifuged at 500 rpm for 1 minute, then left on a plate shaker set at 450 rpm at room temperature for 20 minutes. 20 µL of detection reagent was then added to all wells. The plates were centrifuged at 500 rpm for 1 minute, then left on a plate shaker set at 450 rpm at room temperature for 5 minutes. 10 µL of NaOH was then added to all wells to stop the reaction. The plates were centrifuged at 500 rpm for 1 minute. Plates were then read on a PHERAstar FS (BMG), with absorbance at 610 nm measured.

### Covalent Fragment Intact Protein MS Assay

Recombinantly purified human USP8 catalytic domain (*Hs*USP8_734-1110) (4 µM) in 50 mM HEPES pH 7.5, 150 NaCl was combined with and without compound DDD02155776 (final concentration 100 µM and 1% v/v DMSO) in a final volume of 60 µL. The reaction mixture was incubated at room temperature for 60 mins before quenched by adding 12 µL 1% formic acid (VWR, Radnor, PA).

The LC-MS run were done on an Ultimate 3000 Series high performance liquid chromatogram system (Thermo Scientific, US) coupled with Q-Exactive Orbitrap plus tandem mass spectrometer (Thermo Scientific). Briefly, protein samples were diluted 1 in 2 qith 1% formic acid, and 5 µL diluted sample was initially trapped on an Agilent ZORBAX 300SB-C8 Trap 0.3×5mm 5µm trap and then separated on an Agilent ZORBAX 300SB-C8, 0.075 X 50mm 3.5µm Column. Trap was initially washed with 0.1% TFA at a flow rate of 10 μL/min for 2min before sample progresses to column LC was performed at 50 °C at a flow rate of 1.5 μL/min using mobile phases of 0.1% (v/v) formic acid in water (solvent A) and 0.1% (v/v) formic acid in acetonitrile-water (v/v, 80/20, solvent B). A 35 min gradient was performed with a 3 min of 30% A and the gradient was increased to 35% B by 5 min, followed by an increase to 70% B by 10 min and held for 10 min before returning to the initial conditions. A full MS SIM scan in positive ion mode was obtained for eluting protein in the range of 500-3000 m/z was collected on the Orbitrap at a resolution of 17,500. Raw MS spectra were deconvoluted to neutral monoisotopic masses using the UniDec algorithm (version 4.1.1) (PMID: 25799115) implemented by the Thermo Scientific MSFileReader (version 3.1) with signal-to-noise threshold for peak detection of 5 using charge states less than 60.

### USP8 Aggregation assay

The assay was performed in an assay buffer (pH8 20mM Tris 150mM NaCl 1mM TCEP 10% D2O) in 5mm NMR tubes. The samples are produced from either a solid stock or liquid stock diluted in DMSO to 10mM then diluted further using the assay buffer to 1ml of 200µM. This is then diluted further by serial dilution to produce 500µl samples at concentrations of 200µM, 100µM, 50µM, 25µM and 12.5 µM. These are run on a proton ^1^H experiment on the 500MHz Cryo NMR (Bruker, Avance 3).

The pulse program for the NMR aggregation assay is the standard ^1^H NMR experiment available on all commercial spectrometers. The number of scans was initially 1024 for each experiment changing to a range of scans dependent on concentration (200µM 128 scans, 100µM 256 scans, 50µM 512 scans, 25µM 1024 scans and 12.5µM 1024 scans) to maintain the intensity across all of the experiments. The experiments have a relaxation delay and an acquisition time of 2s, samples were loaded in a sample changer and Icon automation allowed the samples to be queued and run without operator interference.

Data visualisation was one using Bruker’s TOPSPIN software which allows for superposition of 1D NMR spectra along with zooming capabilities. The interpretation of the NMR data was based on an analysis of the superimposed spectra and the observation of major or minor unusual features in resonance shift, shape or number.

### Compound Preparation

#### General chemistry methods

Chemicals and solvents were purchased from commercial vendors and were used as received, unless otherwise stated. Dry solvents were purchased in Sure Seal bottles stored over molecular sieves. Unless otherwise stated herein reactions have not been optimized Analytical thin-layer chromatography (TLC) was performed on precoated TLC plates (Kieselgel 60 F254, BDH). Developed plates were air-dried and analysed under a UV lamp (UV 254/365 nm) and/or KMnO_4_ was used for visualization. Flash chromatography was performed using Combiflash Companion Rf (Teledyne ISCO) and prepacked silica gel columns purchased from Grace Davison Discovery Science or SiliCycle. Mass-directed preparative HPLC separations were performed using a Waters HPLC (2545 binary gradient pumps, 515 HPLC make-up pump, 2767 sample manager) connected to a Waters 2998 photodiode array and a Waters 3100 mass detector. Preparative HPLC separations were performed with a Gilson HPLC (321 pumps, 819 injection module, 215 liquid handler/injector) connected to a Gilson 155 UV/vis detector. On both instruments, HPLC chromatographic separations were conducted using Waters XBridge C18 columns, 19 mm × 100 mm, 5 μm particle size, using 0.1% ammonia in water (solvent A) and acetonitrile (solvent B) as mobile phase.^1^H NMR spectra were recorded on a Bruker Advance II 500 or 400 spectrometer operating at 500 and 400 MHz (unless otherwise stated) using CDCl_3_, DMSO-*d*_*6*_ or CD_3_OD solutions. Chemical shifts (δ) are expressed in ppm recorded using the residual solvent as the internal reference in all cases. Signal splitting patterns are described as singlet (s), doublet (d), triplet (t), multiplet (m), broadened (br) or a combination thereof. Coupling constants (*J*) are quoted to the nearest 0.1 Hertz (Hz). Low resolution electrospray (ES) mass spectra were recorded on a Bruker Daltonics MicrOTOF mass spectrometer run in positive mode. High resolution mass spectroscopy (HRMS) was performed using a Bruker Daltonics MicroTof mass spectrometer. LC−MS analysis and chromatographic separation were conducted with either a Bruker Daltonics MicrOTOF mass spectrometer connected to an Agilent diode array detector or a Thermo Dionex Ultimate 3000 RSLC system with diode array detector, the column used was a Waters XBridge column (50 mm × 2.1 mm, 3.5 μm particle size), and the compounds were eluted with a gradient of 5−95% acetonitrile/water + 0.1% ammonia, or with an Agilent Technologies 1200 series HPLC connected to an Agilent Technologies 6130 quadrupole LC/ MS, connected to an Agilent diode array detector, the column used was a Waters XBridge column (50 mm × 2.1 mm, 3.5 μm particle size) or a Waters X-select column (30 mm × 2.1mm, 2.5 μm particle size) with a gradient of 5-90% acetonitrile/water + 0.1% formic acid, or with an Advion Expression Mass Spectrometer connected to a Thermo Dionex Ultimate 3000 HPLC with diode array detector, the column used was Waters XBridge column (50 mm × 2.1 mm, 3.5 μm particle size) or a Waters X-select column (30 mm × 2.1mm, 2.5 μm particle size) with a gradient of 5-90% acetonitrile/water + 0.1% formic acid. All final compounds showed chemical purity of ≥95% as determined by the UV chromatogram (190−450 nm) obtained by LC−MS analysis. Microwave-assisted chemistry was performed using a CEM or a Biotage microwave synthesizer.

### General method for preparation on ureas 23-25

Potassium carbonate (8.11 mmol) was added to a solution of nitropyridinone (3.24 mmol) in DMF (12 mL) at 0°C followed by alkyl halide (3.89 mmol). The reaction mixture was warmed to RT and stirred overnight. The reaction mixture was partitioned between EtOAc (25 mL) and H_2_O (25 mL) and the layers separated. The aqueous layer was extracted by EtOAc (2 × 25 mL). The combined organics were washed with brine (1 × 25 mL), dried (MgSO4), filtered and concentrated under reduced pressure. Purification using flash column chromatography (Combiflash, 24 g column, 0-50% heptane/EtOAc). Substituted nitropyridinone (0.51 mmol) was taken up in DCM (3 mL) and diluted with Methanol (9 mL). The solution was filtered through a 0.45 uM filter before passing through H-Cube (10% Pd/C CatCart, Full H2, 1 mL/min, RT). The reaction mixture was concentrated in vacuo before taking up in MeOH and passing through 2 g SCX-2. The column was washed with MeOH before the product was eluted with 7M NH3/MeOH. The resultant filtrate was concentrated under reduced pressure to give amino pyridinone, which was used without any further purification in the next step. Amino pyridinone (0.33 mmol) was dissolved in CH_2_Cl_2_ (2 mL) and triethylamine (0.13 mL, 0.90 mmol) added followed by isocyanide (0.54 mmol) and the mixture stirred at room temperature overnight. The reaction mixture was diluted with water (5 mL) and extracting with CH_2_Cl_2_ (2 × 5 mL). The combined organics were passed through a phase separator and concentrated under reduced pressure. The residue was purified using mass directed HPLC (5-95% 0.1% formic acid solution/ acetonitrile) to give ureas **23-25**. 1-isobutyl-3-(1-isopropyl-4-methyl-6-oxo-3-pyridyl)urea **23** ^1^H NMR (500 MHz, DMSO-*d*_*6*_): δ 0.86 (6H, d, *J* = 6.6 Hz), 1.25 (6H, d, *J* = 6.8 Hz), 1.68 (1H, quin, *J* = 6.6 Hz), 2.03 (3H, s), 2.51 (3H, s) 2.88 (2H, m), 3.18 (1H, d, *J* = 5.0 Hz), 5.01 (1H, quin, 6.8 Hz) 6.22 (1H, s), 6.31 (1H, t, *J* = 5.7 Hz) 7.31 (1H, s), 7.68 (1H, s); ^13^C NMR (125 MHz, DMSO-*d*_*6*_): δ 18.0, 20.5, 21.8, 29.0, 45.9, 47.3, 118.4, 120.0, 128.7, 148.2, 157.0, 159.9; HRMS (ESI) calculated for C_14_H_24_N_3_O_2_ [M + H]^+^ 266.1869 found 266.1867; t_R_ = 2.8 min. 1-(1-ethyl-4-methyl-6-oxo-3-pyridyl)-3-isobutyl-urea **24** ^1^H NMR (400 MHz, DMSO-*d*_*6*_): δ 7.67 (1H, s), 7.28 (1H, s), 6.28 (1H, br t), 6.12 (1H, s), 3.85 (2H, q, *J* = 7.1 Hz), 2.87 (2H, t, *J* = 6.2 Hz), 2.02, (3H, s), 1.74 – 1.61 (1H, m), 1.18 (3H, d, *J* = 7.1 Hz) and 0.86 (6H, d, *J* = 6.7 Hz); ^13^C NMR (100 MHz, DMSO-*d*_*6*_) δ 160.1, 157.0, 149.2, 133.3, 119.6, 118.5, 47.3, 43.7, 29.0, 20.5, 18.1 and 15.0; HRMS (ESI) calculated for C_13_H_22_N_3_O_2_ [M + H]^+^ 252.1712 found 252-1725; t_R_ = 2.7 min. 1-(cyclohexylmethyl)-3-(1,4-dimethyl-6-oxo-3-pyridyl)urea **25** ^1^H NMR (400 MHz, DMSO-*d*_*6*_): δ 7.67 (1H, s), 7.25 (1H, s), 6.25 (1H, br t), 6.22 (1H, s), 3.36 (3H, s), 2.89 (2H, t, *J* = 6.3 Hz), 2.02 (3H, s), 1.73 – 1.56 (4H, m), 1.43 – 1.32 (1H, m), 1.26 – 1.09 (4H, m) and 0.97 – 0.80 (2H, m); ^13^C NMR (100 MHz, DMSO-*d*_*6*_) δ 160.3, 157.0, 149.3, 134.6, 119.3, 118.1, 46.0, 38.5, 36.7, 30.8, 26.6, 26.0 and 18.1; HRMS (ESI) calculated for C_15_H_24_N_3_O_2_ [M + H]^+^ 278.1869 found 278.1867; t_R_ = 3.1 min.

### General method for preparation of amides 26-28

Amino pyridinone (0.35 mmol) was dissolved in DMF (2 mL) and triethylamine (1.17 mmol) added followed by 4-methylpentanoic acid (0.56 mmol). T3P (0.70 mmol) was then added and the mixture stirred at room temperature overnight. The reaction mixture was diluted with water (5 mL) and extracting with CH_2_Cl_2_ (2 × 5 mL). The combined organics were passed through a phase separator and concentrated under reduced pressure. The residue was purified using mass directed autoprep (acidic method) to give amides **26-28**. N-(1,4-dimethyl-6-oxo-3-pyridyl)-2-(3-methoxyphenyl)acetamide **26** ^1^H NMR (400 MHz, DMSO-*d*_*6*_): δ 9.29 (1H, br), 7.62 (1H, s), 7.24 (1H, t, *J* = 8.0 Hz), 6.94 – 6.88 (2H, m), 6.85 – 6.80 (1H, m), 6.24 (1H, s), 3.76 (3H, s), 3.57 (2H, s), 3.36 (3H, s) and 1.96 (3H, s); ^13^C NMR (100 MHz, DMSO-*d*_*6*_) δ 170.5, 161.1, 159.7, 149.7, 137.9, 136.5, 129.8, 121.8, 118.1, 117.5, 115.3, 112.5, 55.5, 42.8, 36.7 and 17.9; HRMS (ESI) calculated for C_16_H_19_N_2_O_3_ [M + H]^+^ 287.1396 found 287.1398; t_R_ = 2.8 min. N-(1,4-dimethyl-6-oxo-3-pyridyl)-4-methyl-pentanamide **27** ^1^H NMR (500 MHz, DMSO-*d*_*6*_): δ 9.04 (1H, s), 7.58 (1H, s), 3.37, 2.26 (2H, t, *J* = 7.7 Hz), (3H, s), 1.99 (3H, s), 1.54-1.59 (1H, m), 1.47 (2H, dt, *J* =), 0.89 (6H, d, *J* = 6.6 Hz); ^13^C NMR (125 MHz, DMSO-*d*_*6*_): δ 172.9, 161.1, 149.8, 136.4, 118.0, 117.8, 36.7, 34.7, 33.9, 27.2, 22.7, 18.0; HRMS (ESI) calculated for C_13_H_21_N_2_O_2_ [M + H]+ 237.1603 found 237.1598; t_R_ = 2.8 min. N-[(3-fluorophenyl)methyl]-1-methyl-6-oxo-pyridine-3-carboxamide **28**

^1^H NMR (400 MHz, DMSO-*d*_*6*_): δ 8.76 (1H, br t), 8.39 (1H, d, *J* = 2.6 Hz), 7.89 (1H, dd, *J* = 9.5 and 2.7 Hz), 7.41 - 7.33 (1H, m), 7.18 – 7.03 (3H, m), 6.41 (1H, d, *J* = 9.5 Hz), 4.45 (2H, d, *J* = 5.9 Hz) and 3.48 (3H, s); ^13^C NMR (100 MHz, DMSO-*d*_*6*_) δ 164.1, 162.7 (d, *J*_*C-*F_ = 241.7 Hz), 162.3, 143.1 (d, *J*_*C-*F_ = 6.9 Hz), 142.7, 138.1, 130.6 (d, *J*_*C-*F_ = 8.5 Hz), 123.7 (d, *J*_*C-*F_ = 2.5 Hz), 118.3, 114.4 (d, *J*_*C-*F_ = 21.4 Hz), 113.9 (d, *J*_*C-*F_ = 20.8 Hz), 112.5, 42.5 and 37.7; HRMS (ESI) calculated for C_14_H_14_FN_2_O_2_ [M + H]^+^ 269.1039 found 261.1044; t_R_ = 2.9 min.

### General method for preparation of amides 29-30

1-Methyl-2-pyrrolecarboxylic acid (4.00 mmol) was dissolved in CH_2_Cl_2_ (10 mL) and aluminium chloride (8.00 mmol) added in an ice bath. acid chloride (4.40 mmol) was added while the mixture was in the ice bath. The reaction was then heated at 45oC overnight. The mixture was added to saturated sodium bicarbonate solution (20 mL) and the mixture filtered through celite. The aqueous layer was acidified with conc. HCl and the precipitate collected by filtration to give acid intrermediate.

Acid intermediate (0.60 mmol) was dissolved in CH_2_Cl_2_ (3 mL) and benzylamine (0.72 mmol) added followed by triethylamine (0.90 mmol). T3P in EtOAc (0.72 mmol) was added and the mixture stirred overnight. Saturated sodium bicarbonate solution (2 mL) was added to the mixture, which was stirred for 1 hour. The organic layer was collected and the solvent removed. The residue was purified by flash chromatography, gradient elution from heptane to EtOAc, to give amides **29** and **30**.

N-benzyl-4-(2-chloroacetyl)-1-methyl-pyrrole-2-carboxamide **29**

^1^H NMR (500 MHz, CDCl_3_): δ 7.40 (1H, d, *J* = 1.6 Hz), 7.25-7.37 (5H, m), 7.03, (1H, d, *J* = 1.6 Hz), 6.39,

(1H, br s), 4.57 (2H, d, *J* = 5.8 Hz), 4.35 (2H, s), 3.99 (3H, s); ^13^C NMR (125 MHz, CDCl_3_) δ 186.1, 160.8, 138.0, 131.8, 128.8, 127.8, 127.7, 120.5, 111.8, 25.7, 43.5, 37.6; HRMS (ESI) calculated for C_15_H_16_N_2_O_2_Cl [M + H]^+^ 291.0895 found 291.0915; t_R_ = 3.3 min.

4-acetyl-N-benzyl-1-methyl-pyrrole-2-carboxamide **30**

^1^H NMR (500 MHz, CDCl_3_): δ 7.29-7.38 (6H, m), 6.97, (1H, d, *J* = 1.8 Hz), 6.28, (1H, br s), 4.58 (2H, d, *J* = 5.8 Hz), 4.00 (3H, s), 2.38 (3H, s); ^13^C NMR (125 MHz, CDCl_3_) δ 192.6, 161.1, 138.1, 131.4, 128.8, 127.8, 127.7, 127.3, 124.2, 111.4, 43.5, 37.4, 27.0; HRMS (ESI) calculated for C_15_H_17_N_2_O_2_ [M + H]^+^ 257.1285 found 257.1304; t_R_ = 3.0 min.

### General method for preparation of compounds 33-37

To a solution of quinoxaline-2-carboxylate ester (0.23 mmol) in Ethanol (1.5 mL) was added amine (0.34 mmol) and then triethylamine (0.08 mL, 0.57 mmol). The reaction was heated in the microwave at 120 °C for 30 mins. The reaction mixture was concentrated under reduced pressure and purified using mass directed autoprep (basic method) to give amides **33**-**37**.

N,4,6,7-tetramethyl-3-oxo-quinoxaline-2-carboxamide **33**

^1^H NMR (500 MHz, DMSO-*d*_*6*_): δ 8.78 (1H, br), 7.65 (1H, s), 7.46 (1H, s), 3.64 (3H, s), 2.80 (3H, d, *J* = 4.8 Hz), 2.43 (3H, s) and 2.34 (3H, s); ^13^C NMR (125 MHz, DMSO-*d*_*6*_) δ 164.3, 153.6, 149.7, 142.5, 133.3, 132.2, 130.4, 130.2, 115.7, 29.5, 26.1, 20.6 and 19.1; LCMS 245.9 @ 1.2 min. 6,7-dimethyl-N-(3-morpholinopropyl)-3-oxo-4H-quinoxaline-2-carboxamide **34**

^1^H NMR (400 MHz, DMSO-*d*_*6*_): δ 12.41 (1H, br), 9.14 (1H, s), 7.62 (1H, s), 7.14 (1H, s), 3.58 (4H, t, *J* = 4.6 Hz), 3.36 – 3.30 (2H, m, under H_2_O peak), 2.42 – 2.28 (12H, m) and 1.68 (2H, qn, *J* = 6.9 Hz); ^13^C NMR (100 MHz, DMSO-*d*_*6*_) δ 164.4, 154.9, 142.3, 133.2, 131.5, 130.5, 129.4, 117.3, 116.2, 66.7, 56.4, 53.8, 37.7, 26.2, 20.4 and 19.4; HRMS (ESI) calculated for C_18_H_25_N_4_O_3_ [M + H]^+^ 345.1927 found 345.1915;

LCMS 345.0 @ 1.0 min.

N-(2-methoxyethyl)-3-oxo-3,4-dihydroquinoxaline-2-carboxamide **35**

^1^H NMR (400 MHz, DMSO-*d*_*6*_): δ 12.8 (1H, br), 9.13 (1H, br), 7.86 (1H, d, *J* = 7.3 Hz), 7.63 (1H, t, *J* = 7.0 Hz), 7.41 – 7.34 (2H, m), 3.52 – 3.42 (4H, m) and 3.30 (3H, s); ^13^C NMR (100 MHz, DMSO-*d*_*6*_) d 163.4, 154.5, 151.3, 132.9, 132.3, 131.8, 129.8, 124.3, 116.0, 70.8, 58.5 and 39.1; HRMS (ESI) calculated for C_12_H_14_N_3_O_3_ [M + H]^+^ 248.1035 found 248.1025; LCMS 248.0 @ 1.0 min.

Ethyl 7-methyl-3-oxo-3,4-dihydroquinoxaline-2-carboxylate **36**

^1^H NMR (400 MHz, DMSO-*d*_*6*_): δ 12.70 (1H, br), 8.90 (1H, br), 7.65 (1H, s), 7.46 (1H, d, *J* = 8.3 Hz), 7.26 (1H, d, *J* = 8.3 Hz), 2.82 (3H, d, *J* = 4.8 Hz) and 2.40 (3H, s); ^13^C NMR (100 MHz, DMSO-*d*_*6*_) δ 164.0, 154.3, 151.7, 133.7, 133.5, 131.7, 130.7, 129.1, 115.7, 26.1 and 20.8; HRMS (ESI) calculated for C_11_H_12_N_3_O_2_ [M + H]^+^ 218.0930 found 218.0919; LCMS 217.9 @ 1.1 min.

N,N-dimethyl-3-oxo-3,4-dihydroquinoxaline-2-carboxamide **37**

^1^H NMR (500 MHz, DMSO-*d6*): δ 12.77 (1H, s), 7.81 – 7.76 (1H, m), 7.62 – 7.57 (1H, m), 7.38 – 7.32 (2H, m), 3.00 (3H, s) and 2.90 (3H, s); ^13^C NMR (120 MHz, DMSO-*d6*) δ 165.23, 155.8, 153.2, 132.7, 131.6, 129.2, 124.2, 116.2, 37.3 and 34.1; LCMS 218.0 @ 1.0 min; HRMS (ESI) calculated for C_11_H_11_N_3_O_2_ [M + H]^+^ 218.0930 found 218.1264

3-(4-benzoylpiperazin-1-yl)-5,6-dimethyl-pyridazine-4-carbonitrile **39**

3-chloro-5,6-dimethyl-pyridazine-4-carbonitrile (5.97mmol) was dissolved in DMSO (30mL) and piperazine (29.83mmol) added. The mixture was heated for 1 hour at 130 °C in the microwave. The solvent was removed, and the residue was diluted in DCM (20mL) and washed with sat. NaHCO3 solution (20mL). The residue was purified using a SCX column to give 5,6-dimethyl-3-piperazin-1-yl-pyridazine-4-carbonitrile.

5,6-Dimethyl-3-piperazin-1-yl-pyridazine-4-carbonitrile (0.46 mmol) and potassium carbonate (127 mg,0.92 mmol) were suspended in MeCN (3mL) and benzoyl chloride (0.059 mL, 0.51 mmol) added. The mixture was stirred overnight. The mixture was purified directly by mass directed autoprep (basic method) to give 3-(4-benzoylpiperazin-1-yl)-5,6-dimethyl-pyridazine-4-carbonitrile.

^1^H NMR (500 MHz, DMSO-*d*_*6*_): δ 7.46 (br s, 5H), 3.57-3.77 (8H, m), 2.55 (3H, s), 2.43 (3H, s); ^13^C NMR (125 MHz, DMSO-*d*_*6*_) δ 169.7, 159.3, 154.3, 143.2, 136.2, 130.1, 128.9, 127.5, 115.2, 104.2, 49.2, 49.1, 47.3, 41.8, 20.0, 17.9; HRMS (ESI) calculated for C_18_H_20_N_5_O [M + H]^+^ 322.1662 found 322.1657; t_R_ = 3.3 min.

1-(4-cyano-5,6-dimethyl-pyridazin-3-yl)-N-methyl-piperidine-4-carboxamide **40**

3-chloro-5,6-dimethylpyridazine-4-carbonitrile (0.6 mmol) was dissolved in DMSO (3 mL) and Triethylamine (0.9 mmol) and n-methylpiperidine-4-carboxamide (0.66 mmol) added. The mixture was heated for 1 hour at 130 °C in the microwave. The mixture was purified by mass directed autoprep (basic method) to give 1-(4-cyano-5,6-dimethyl-pyridazin-3-yl)-N-methyl-piperidine-4-carboxamide. ^1^H NMR (500 MHz, DMSO-*d*_*6*_): δ 7.76 (1H, s), 4.04 (2H, d, *J* = 13.1 Hz), 3.03 (2H, t, *J* = 13.1 Hz), 2.59 (3H, d, *J* = 4.5 Hz), 2.53 (3H, s), 2.41 (3H, s), 2.40 – 2.34 (1H, m) and 1.85 – 1.66 (4H, m); ^13^C NMR (125 MHz, DMSO-*d*_*6*_) δ 174.7, 159.7, 153.5, 142.9, 115.3, 103.8, 49.0, 41.9, 28.6, 26.0, 19.9 and 17.8; HRMS (ESI) calculated for C_14_H_20_N_5_O [M + H]^+^ 274.1668 found 274.1686; t_R_ = 3.3 min.

### General procedure for scavenging metal contaminants

Compound (0.13 mmol) was dissolved in Methanol (5 mL) and MP-TMT (0.13 mmol) added. The mixture was stirred at room temperature overnight. The polymer scavenger was removed by filtration through a TMT scavenger column to give scavenged material.

3-chloro-5,6-dimethyl-pyridazine-4-carbonitrile **41**

^1^H NMR (500 MHz, DMSO-*d*_*6*_): δ 2.66 (3H, s) and 2.52 (3H, s); ^13^C NMR (125 MHz, DMSO-*d*_*6*_) δ 160.9, 152.5, 145.2, 114.6, 113.4, 20.4 and 18.5.

2-tert-butyl-5-pyrrolidin-1-yl-1,3,4-thiadiazole **4**2

^1^H NMR (500 MHz, MeOD): δ 3.44-3.47 (4H, m), 2.05-2.08 (4H, m), 1.30 (9H, s); ^13^C NMR (125 MHz, MeOD) δ 168.6, 168.5, 50.3, 35.6, 29.6, 25.3; LCMS 211.9 @ 2.7 min.

2-tert-butyl-5-(1-piperidyl)-1,3,4-thiadiazole **43**

^1^H NMR (500 MHz, MeOD): δ 3.46-3.47 (4H, m), 1.68 (6H, br s), 1.39 (9H, s); ^13^C NMR (125 MHz, MeOD): δ 171.0, 168.4, 49.3, 34.2, 28.1, 23.3, 22.1; LCMS 225.9 @ 3.5 min.

